# Inferring tree causal models of cancer progression with probability raising

**DOI:** 10.1101/000919

**Authors:** Loes Olde Loohuis, Giulio Caravagna, Alex Graudenzi, Daniele Ramazzotti, Giancarlo Mauri, Marco Antoniotti, Bud Mishra

## Abstract

Existing techniques to reconstruct tree models of progression for accumulative processes, such as cancer, seek to estimate causation by combining correlation and a frequentist notion of temporal priority. In this paper, we define a novel theoretical framework called CAPRESE (CAncer PRogression Extraction with Single Edges) to reconstruct such models based on the notion of probabilistic causation defined by Suppes. We consider a general reconstruction setting complicated by the presence of noise in the data due to biological variation, as well as experimental or measurement errors. To improve tolerance to noise we define and use a shrinkage-like estimator. We prove the correctness of our algorithm by showing asymptotic convergence to the correct tree under mild constraints on the level of noise. Moreover, on synthetic data, we show that our approach outperforms the state-of-the-art, that it is efficient even with a relatively small number of samples and that its performance quickly converges to its asymptote as the number of samples increases. For real cancer datasets obtained with different technologies, we highlight biologically significant differences in the progressions inferred with respect to other competing techniques and we also show how to validate conjectured biological relations with progression models.

## Introduction

Cancer is a disease of evolution. Its initiation and progression are caused by dynamic somatic alterations to the genome manifested as point mutations, structural alterations, DNA methylation and histone modification changes [1].

These genomic alterations are generated by random processes, and since individual tumor cells compete for space and resources, the fittest variants are naturally selected for. For example, if through some mutations a cell acquires the ability to ignore anti-growth signals from the body, this cell may thrive and divide, and its progeny may eventually dominate some part(s) of the tumor. This *clonal expansion* can be seen as a *discrete state* of the cancer’s progression, marked by the acquisition of a set of genetic events. Cancer progression can then be thought of as a sequence of these discrete steps, where the tumor acquires certain distinct properties at each state. Different progression sequences are possible, but some are more common than others, and not every order is viable [2].

In the last two decades, many specific genes and genetic mechanisms that are involved in different types of cancer have been identified (see e.g. [3, 4] for an overview of common cancer genes and [5, 6] for specific genetic analyses of ovarian carcinoma and lung adenocarcinoma, respectively), and *therapies* targeting the activity of these genes are now being developed at a fast pace [2]. However, unfortunately, the *causal and temporal relations* among the genetic events driving cancer progression remain largely elusive.

The main reason for this state of affairs is that information revealed in the data is usually obtained only at one (or a few) points in time, rather than over the course of the disease. Extracting this dynamic information from the available *cross-sectional* data is challenging, and a combination of mathematical, statistical and computational techniques is needed. In recent years, several methods to extract progression models from cross-sectional data have been developed, starting from the seminal work on single-path-models by Fearon and Vogelstein [7]. In particular, different models of oncogenetic trees were developed over the years. At the core of some of these methods, e.g. [8, 9], is the use of *correlation* to identify relations among genetic events. These techniques reconstruct *tree* models of progression as independent acyclic paths with branches and no confluences. Distinct models of oncogenetic trees are instead based on *maximum likelihood estimation,* e.g., [10, 11, 12]. More general *Markov chain* models, e.g., [13], describe more flexible probabilistic networks, despite the computationally expensive parameter estimation. Other recent models are Conjunctive Bayesian Networks, CBNs [14, 15], that extract *directed acyclic graphs,* yet imposing specific constraints on the joint occurrence of events. Finally, in a slightly different context, temporal models were reconstructed from time-course gene expression data [16, 17].

In this paper we present a novel theoretical framework called CAPRESE (CAncer PRogression Extraction with Single Edges) to reconstruct cumulative progressive phenomena, such as cancer progression. We assume the original problem setting of [8], and propose a new a technique to infer *probabilistic progression trees* from cross-sectional data. Unlike maximum likelihood estimation-based techniques, our goal is the extraction of the *minimal* progression model explaining the order in which mutations occur and accumulate. The method is technology agnostic, i.e., it can be applied to dataset derived from all types of (epi-)genetic data such as deep exome sequencing, bisulfite sequencing, SNP arrays, etc., (see Results), and takes as input a set of pre-selected genetic events of which the presence or the absence of each event is recorded for each sample.

CAPRESE is based on two main ingredients: 1) rather than using *correlation* to infer progression structures, we base our technique on a notion of *probabilistic causation,* and 2), to increase robustness against noise, we adopt a *shrinkage-like estimator* to measure causation among any pair of events.

More specifically, with respect to our first ingredient, we adopt the notion of (prima facie) causation proposed by Suppes in [18]. Its basic intuition is simple: event *a* causes event *b* if (*i*) *a* occurs *before b* and (*ii*) the occurrence of *a raises the probability* of observing b. This is a very basic notion of probabilistic causation that in itself does not address many of the problems associated with it (such as asymmetry, common causes, and screening off [19]), and includes *spurious* as well as *genuine* causes. However, as it turns out, this basic notion combined with a filter for independent progressions starting from the same root, is an excellent tool to guide progression extraction from cross-sectional data - one that outperforms the commonly used correlation-based methods.

Probabilistic causation was used in biomedical applications before (e.g., to find driver genes from CNV data in [20], and to extract causes from biological time series data in [21]), but, to the best of our knowledge, never to infer *progression models* in the *absence* of direct temporal information.

The extraction problem is complicated by the presence of both false positive and false negative observations (see [22] for a discussion on this issue based on the reconstruction by [8]), such as the one provided by the intrinsic variability of biological processes (e.g., *genetic heterogeneity)* and *experimental errors.* This poses a problem, because while probability raising is a very precise tool, it, by itself, is not robust enough against noise. Conditional on the amount of noise, we will rely both on probabilistic causation and on a more robust (but less precise) correlation-based metric in an optimal way. To do this we introduce our second ingredient, a *shrinkage-like estimator* to measure causation among any pair of events. The intuition behind this estimator, which is closely related to a shrinkage estimator from [23], is to find the optimal balance between probability raising on the one hand and correlation on the other, depending on the amount of noise.

We prove correctness of our algorithm by showing that with increasing sample sizes, the reconstructed tree asymptotically converges to the correct one (Theorem 3). Under mild constraints on the noise rates, this result holds for the reconstruction problem in the presence of uniform noise as well.

We also study the performance of CAPRESE in more realistic settings with limited sample sizes. Using synthetic data, we show that under these conditions, our algorithm outperforms the state-of-the-art tree reconstruction algorithm of [8] (see Results). In particular, our shrinkage-like estimator provides, on average, an increased robustness to noise which ensures it to outperform oncotrees [8]. Performance is defined in terms of *structural similarity* between the reconstructed tree and the actual tree, rather than on their induced distribution as is done, e.g., in [11]. This metric is especially appropriate for the goal of reconstructing a progression model where data-likelihood fit is secondary to “calling” the possibly minimal set of causal relations.

Also, we show that CAPRESE works well already with a relatively low number of samples and that its performance quickly converges to its asymptote as the number of samples increases. This outcome hints at the applicability of the algorithm with relatively small datasets without compromising its efficiency.

We remark that further analyses on synthetic data suggests that CAPRESE outperforms a well known bayesian probabilistic graphical model as well (i.e., *Conjunctive Bayesian Networks* [14, 15]), which was originally conceived for the reconstruction of more complex topologies, e.g. DAGs, but was proven effective in reconstructing tree topologies as well [24] (see Results).

Finally, we apply our technique to alterations assessed with both Comparative Genomic Hybridization and Next Generation Sequencing techniques (see Results). In the former case, we show that the algorithm of [8] and CAPRESE highlight biologically important differences in ovarian, gastrointestinal and oral cancer, but our inferences are statistically more significant. In the latter, we validate a recently discovered relation among two key genes involved in leukemia.

## Methods

### Problem Setting

The set-up of the reconstruction problem is as follows. Assuming that we have a set *G* of *n* mutations (*events,* in probabilistic terminology) and s samples, we represent a cross-sectional dataset as an s × n binary matrix in which an entry (*k, l*) = 1 if the mutation *l* was observed in sample *k,* and 0 otherwise. The problem we solve in this paper is to extract a set of edges *E* yielding a progression *tree* 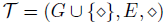 from this matrix which, we remark, only implicitly provides information of progression timing. The root of *𝒯* is modeled using a (special) event 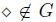 *heterogenous progression paths* or *forests* can be reconstructed. More precisely, we aim at reconstructing a *rooted tree* that satisfies: (*i*) each node has at most one incoming edge, (*ii*) the root has no incoming edges (*iii*) there are no *cycles*.

Each progression tree subsumes a distribution of observing a subset of the mutations in a cancer sample that can be formalized as follows:

**Definition 1** (Tree-induced distribution). *Let 𝒯 be a tree and α : E→ [0,1] a labeling function denoting the independent probability of each edge, T generates a distribution where the probability of observing a sample with the set of alterations G*^*^ ⊆ *G* is 
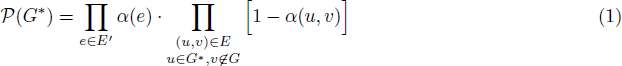
where all events in *G*^*^ are assumed to be reachable from the root ◊, and *E*′ ⊆ *E is the set of edges connecting the root to the events in G^*^*.

We would like to emphasize two properties related to the tree-induced distribution. First, the distribution subsumes that, given any oriented edge *(a → b),* an observed sample contains alteration *b* with probability 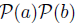, that is the probability of observing *b* after *a*. For this reason, if *a* causes b, the probability of observing *a* will be greater than the probability of observing *b* accordingly to the temporal priority principle which states that all causes must precede, in time, their effects [25].

Second, the input dataset is a set of samples generated, ideally, from an unknown distribution induced by an unknown tree or forest that we aim at reconstructing. However, in some cases, it could be that no tree exists whose induced distribution generates *exactly* those input data. When this happens, the set of observed samples slightly diverges from any tree-induced distribution. To model these situations a notion of *noise* can be introduced, which depends on the context in which data are gathered. Adding noise to the model complicates the reconstruction problem (see Results).

**The *oncotree* approach**. In [8] Desper *et al*. developed a method to extract progression trees, named *“oncotrees”,* from static CNV data. In [22] Szabo *et al*. extended the setting of Desper’s reconstruction problem to account for both *false positives* and *negatives* in the input data. In these oncotrees, nodes represent CNV events and edges correspond to possible progressions from one event to the next.

The reconstruction problem is exactly as described above, and each tree is rooted in the special event ◊. The choice of which edge to include in a tree is based on the estimator 
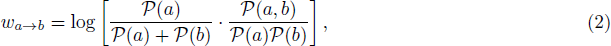
which assigns to each edge *a →b* a weight accounting for both the relative and joint frequencies of the events - thus measuring *correlation*. The estimator is evaluated after including ◊ to each sample of the dataset. In this definition the rightmost term is the (symmetric) *likelihood ratio* for *a* and *b* occurring together, while the leftmost is the asymmetric *temporal priority* measured by rate of occurrence. This implicit form of timing assumes that, if a occurs *more often* than *b*, then it likely occurs *earlier*, thus satisfying 
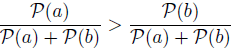

An oncotree is the rooted tree whose total weight (i.e., sum of all the weights of the edges) is maximized, and can be reconstructed in 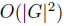 steps using Edmond’s algorithm [26]. By construction, the resulting graph is a proper tree rooted in ◊: each event occurs only once, *confluences* are absent, i.e., any event is caused by at most one other event. This method has been used to derive progressions for various cancer datasets e.g., [27, 28, 29]), and even though several methods that extend this framework exists (e.g. [9, 11, 15]), to the best of our knowledge, it is currently the only method that aims to solve exactly the same problem as the one investigated in this paper and thus provide a benchmark to compare against.

### A Probabilistic Approach to Causation

We briefly review the approach to probabilistic causation, on which our method is based. For an extensive discussion on this topic we refer to [19].

In his seminal work [18], Suppes proposed the following notion.

**Definition 2** (Probabilistic causation, [18]). *For any two events c* and *e*, *occurring respectively at times t_c_ and t_e_, under the mild assumptions that* 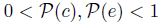, *the event c is a* prima facie cause *of the event e if it occurs* before *the effect and the cause* raises the probability *of the effect, i.e.,*

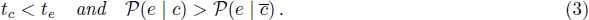

As discussed in [19] the above conditions are not, in general, sufficient to claim that event *c* is a cause of event *e*. In fact a prima facie cause is either *genuine* or *spurious*. In the latter case, the fact that the conditions hold in the observations is due either to coincidence or to the presence of a certain third *confounding factor,* related both to *c* and to *e* [18]. Genuine causes, instead, satisfy Definition 2 and are not screened-off by any confounding factor. However, they need not be direct causes. See Figure 1.

**Figure 1:**
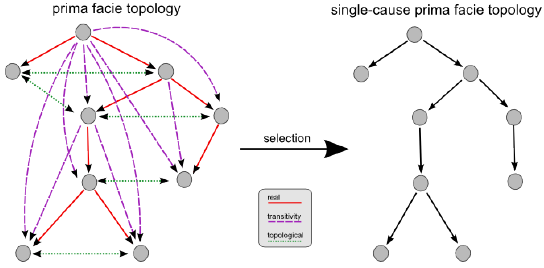
Prima facie topology. Example prima facie topology where all edges (*a*, *b*) represent prima facie causes, according to Definition 3: *a* is a probability raiser of b and it occurs more frequently. In left, we filter out spurious causes and select only the real ones among the genuine, yielding a single-cause prima facie topology.

Note that weconsider cross-sectional data where no information about *t_c_* and *t_e_* is available, so in our reconstruction setting we are restricted to consider solelythe *probability raising* (pr) property, i.e., 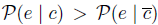, which makes it harder to discriminate among genuine and spurious causes. Now we review some of its properties.

**Proposition 1** (Dependency). *Whenever the* pr *holds between two events a and b, then the events are* statistically dependent *in a positive sense*, i.e.,

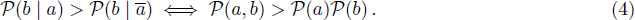

This and the next proposition are well-known facts of the PR; their derivation as well as the proofs of all the results we present is in the Supplementary Materials. Notice that the opposite implication holds as well: when the events *a* and *b* are still dependent but in a negative sense, i.e., 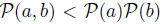, the PR does not hold, i.e., 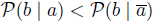.

We would like to use the asymmetry of the PR to determine whether a pair of events *a* and *b* satisfy a causation relation so to place *a* before *b* in the progression tree but, unfortunately, the PR satisfies the following property.

**Proposition 2** (Mutual PR). 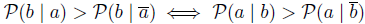.

That is, if *a* raises the probability of observing *b*, then *b* raises the probability of observing *a* too.

Nevertheless, in order to determine causes and effects among the genetic events, we can use our *degree of confidence* in our estimate of probability raising to decide the direction of the causation relationship between pairs of events. In other words, if a raises the probability of *b more* than the other way around, then a is a more likely cause of *b* than *b* of *a*. Notice that this is sound as long as each event has *at most* one cause; otherwise, *frequent late events* with more than one cause, which are rather common in biological progressive phenomena, should be treated differently. As mentioned, the pr is not symmetric, and the *direction* of probability raising depends on the relative frequencies of the events. We make this asymmetry precise in the following proposition.

**Proposition 3** (Probability raising and temporal priority). *For any two events a and b such that the probability raising* 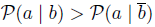 *holds, we have*

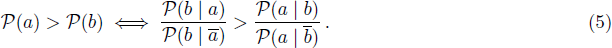

That is, given that the pr holds between two events, a raises the probability of b *more* than *b* raises the probability of *a*, if and only if a is observed more frequently than *b*. Notice that we use the ratio to assess the pr inequality. The proof of this proposition is technical and can be found in the Supplementary Materials. From this result it follows that if we measure the timing of an event by the rate of its occurrence (that is, 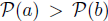 implies that a happens before *b*), this notion of pr subsumes the same notion of temporal priority induced by a tree. We also remark that this is also the temporal priority made explicit in the coefficients of Desper’s method. Given these results, we define the following notion of causation.

**Definition 3**. *We state that a is a* prima facie cause *of b if a is a probability raiser of b, and it occurs more frequently*: 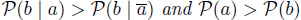

We term *prima facie topology* a directed acyclic graph (over some events) where each edge represents a prima facie cause. When at most a single incoming edge is assigned to each event (i.e., an event has at most a *unique cause,* in the real world), we term this structure *single-cause prima facie topology*. Intuitively, this last class of topologies correspond to the trees or, more generally forests when they have disconnected components, that we aim at reconstructing.

Before moving on to introducing our algorithm let us discuss our definition of *causation*, its role in the definition of the reconstruction problem and some of its limitations. As already mentioned, it may be that for some prima facie cause *c* of an event *e*, there is a third event *a* prior to both, such that *a* causes *c* and ultimately *c* causes *e*. Alternatively, *a* may cause both *c* and *e* independently, and the causation relationship observed from *c* to *e* is merely *spurious*. In the context of the tree-reconstruction problem, namely when it is assumed that each event has at most a unique cause, the aim is to filter out the spurious edges from a general prima facie topology, so to extract a single-cause prima facie structure (see Figure 1).

Definition 3 summarizes Suppes basic notion of prima facie cause, while it is ignoring deeper discussions of causation that aim at distinguishing between actual genuine and spurious causes, e.g. screening-off, background context, d-separation [30, 31, 19]. For our purposes however, the above definition is sufficient when (*i*) all the significant events are considered, i.e., all the genuine causes are observed as in a closed-world assumption, and (*ii*) we aim at extracting the *order* of progression among them (or determine that there is no apparent relation), rather than extracting causalities *per se*. Note that these assumptions are strong and might be weakened in the future (see Discussions), but are shared by us and [8].

Finally, we recall a few algebraic requirements necessary for our framework to be well-defined. First of all, the PR must be computable: every mutation *a* should be observed with probability strictly 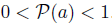. Moreover, we need each pair of mutations (*a, b)* to be *distinguishable* in terms of PR, that is, for each pair of mutations *a* and *b*, 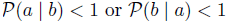 similarly to the above condition. Any non-distinguishable pair of events can be merged as a single composite event. From now on, we will assume these conditions to be verified.

### Performance Measure and Synthetic Datasets

We made use of *synthetic data* to evaluate the performance of CAPRESE as a function of dataset size and the false positive and negative rates. Many distinct synthetic datasets were created for this purpose, as explained below. The algorithm’s performance was measured in terms of *Tree Edit Distance* (TED, [32]), i.e., the minimum-cost sequence of node edit operations (relabeling, deletion and insertion) that transforms the reconstructed trees into the ones generating the data. The choice of this measure of evaluation is motivated by the fact that we are interested in the *structure* behind the progressive phenomenon of cancer evolution and, in particular, we are interested in a measure of the genuine causes that we miss and of the spurious causes that we fail to recognize (and eliminate). Also, since topologies with similar distributions can be structurally different we choose to measure performance using structural distance rather than a distance in terms of distributions. Within the realm of ‘structural metrics’ however, we have also evaluated the performance with the *Hamming Distance* [33], another commonly used structural metric, and we obtained analogous results (not shown here).

**Synthetic data generation and experimental setting**. Synthetic datasets were generated by sampling from various random trees constrained to have depth 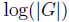, since wide branches are harder to reconstruct than straight paths, and by sampling event probabilities in [0.05,0.95] (see Supplementary Materials).

Unless explicitly specified, in all the experiments we used 100 distinct random trees (or forests, accordingly to the test to perform) of 20 events each. This seems a fairly reasonable number of events and is in line with the usual size of reconstructed trees, e.g. [34, 35, 36, 37]. The *scalability* of the techniques was tested against the number of samples by ranging 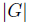 from 50 to 250, with a step of 50, and by replicating 10 independent datasets for each parameters setting (see the caption of the figures for details).

We included a form of *noise* in generating the datasets, in order to account for (i) the realistic presence of *biological noise* (such as the one provided by bystander mutations, genetic heterogeneity, etc.) and (ii) *experimental errors*. A noise parameter 0 ≤ *v* <1 denotes the probability that any event assumes a random value (with uniform probability), after sampling from the tree-induced distribution. Algorithmically this process generates, on average, 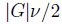 random entries in each sample (e.g. with *v* = 0.1 we have, on average, one error per sample). We wish to assess whether these noisy samples can mislead the reconstruction process, even for low values of *v*. Notice that assuming a uniformly distributed noise may appear simplistic since some events may be more robust, or easy to measure, than others. However, introducing in the data both *false positives* (at rate 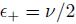 and *negatives* (at rate 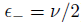 makes the inference problem substantially harder, and was first investigated in [22].

In the Results section, we refer to datasets generated with rate *v* >;0 as noisy synthetic dataset. In the numerical experiments, *v* is usually discretized by 0.025, (i.e., 2.5% noise).

## Results

### Extracting Progression Trees With Probability Raising and A Shrinkage-Like Estimator

The CAPRESE reconstruction method is described in Algorithm 1. The algorithm is similar to Desper’s and Szabo’s algorithm, the main difference being an alternative weight function based on a shrinkage-like estimator.

**Algorithm 1.**
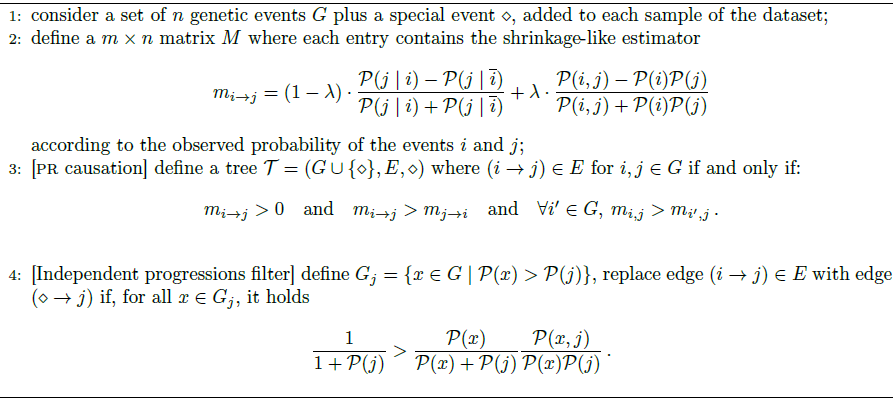
CAPRESE: a tree-like reconstruction with a shrinkage-like estimator

**Definition 4** (Shrinkage-like estimator). *We define the* shrinkage-like estimator *m_a→b_ of the confidence in the causation relationship from a to b as*

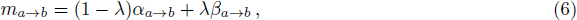
*where* ≤ λ ≤ 1 *and* 
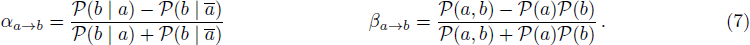

This estimator is similar in spirit to a shrinkage estimator (see [23]) and combines a normalized version of PR, the *raw estimate a*, with a *correction factor* β (in our case a correlation-based measure of temporal distance among events), to define a proper order in the confidence of each causation relationship. Our A is the analogous of the *shrinkage coefficient* and can have a Bayesian interpretation based on the strength of our belief that *a* and *b* are causally relevant to one another and the evidence that *a* raises the probability of *b*. In the absence of a closed form solution for the optimal value of α, one may rely on cross-validation of simulated data. The power of shrinkage (and our shrinkage-like estimator) lies in the possibility of determining an optimal value for α to balance the effect of the correction factor on the raw model estimate to ensure optimal performances on ill posed instances of the inference problem. A crucial difference, however, between our estimator and classical shrinkage, is that our estimator aims at improving the performance of the *overall* reconstruction process, not limited to the performance of the estimator itself as is the case in shrinkage. That is, the metric *m* induces an ordering to the events reflecting our confidence for their causation. Furthermore, since we make no assumption about the underlying distribution, we learn it empirically by cross-validation. In the next sections we show that the shrinkage-like estimator is an effective way to get such an ordering especially when data are noisy. In CAPRESE we use a pairwise matrix version of the estimator.

**The raw estimator and the correction factor**. By considering only the raw estimator α, we would include an edge (*a → b*) in the tree consistently in terms of (*i*) Definition 3 (Methods) and (*ii*) if a is the best probability raiser for *b*. When 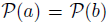 the events *a* and *b* are indistinguishable in terms of temporal priority, thus α is not sufficient to decide their causal relation, if any. This intrinsic ambiguity is unlikely in practice even if, in principle, it is possible. Notice that this formulation of α is a monotonic normalized version of the PR ratio.

**Proposition 4** (Monotonic normalization). *For any two events a and b we have* 
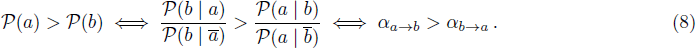

This raw model estimator satisfies 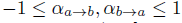: when it tends to —1 the pair of events appear disjointly (i.e., they show an anti-causation pattern), when it tends to 0 no causation or anti-causation can be inferred and the two events are statistically independent and, when it tends to 1, the causation relationship between the two events is genuine. Therefore, a provides a quantification of the degree of confidence for a pr causation relationship. In fact, for any given possible causation edge (*a, b*), the term 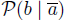 gives an estimate of the *error rate* of b, therefore the numerator of the raw model a provides an estimate of how often b is actually caused by *a*. The a estimator is then normalized to range between —1 and +1.

However, α does not provide a general criterion to disambiguate among genuine causes of a given event. We show a specific case in which a is not a sufficient estimator. Let us consider, for instance, a causal linear path: *a* → *b* → *c*. In this case, when evaluating the candidate parents *a* and *b* for *c* we have: *α_α→c_ = α_b→c_ = 1*, so *a* and *b* are genuine causes of *c*, though we would like to select *b*, instead of *a*. Accordingly, we can only infer that *t_a_<t_c_* and *t_b_<t_c_*, i.e., a partial ordering, which does not help to disentangle the relation among *a* and *b* with respect to *c*.

In this case, the β coefficient can be used to determine which of the two genuine causes occurs closer, in time, to *c* (*b*, in the example above). In general, such a correction factor provides information on the *temporal distance* between events, in terms of statistical dependency. In other words, the higher the β coefficient, the closer two events are. Therefore, when dealing with noisy data and limited sample sizes, there are situations where, by using the α estimator alone, we could infer a wrong transitive edge to be the most likely cause even in the presence of the real cause. For this reason, we reduce the a estimator to the correction factor β, which, for each given edge (*a, b*), is normalized within —1 and 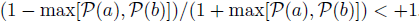.

The shrinkage-like estimator *m* then results in the combination of the raw PR estimator α and of the β correction factor, which respects the temporal priority induced by α.

**Proposition 5** (Coherence in dependency and temporal priority). *The β correction factor is* symmetrical *and subsumes the same notion of dependency of the raw estimator α, that is*

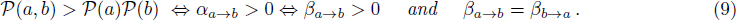

The independent progressions filter. As in Desper’s approach, we also add a *root* ◊ with 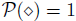 in order to separate different progression paths, i.e., the different sub-trees rooted in ◊. CAPRESE initially builds a unique tree by using the estimator; typically, the most likely event will be at the top of the progression even if there may be rare cases where more than one event has no valid parent, in these cases we would already be reconstructing a forest. In the reconstructed tree, all the edges represent the most confident prima facie cause, although some of those could still be spurious causes. Then the correlation-like weight between any node *j* and ◊ is computed as 
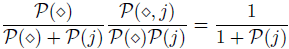

If this quantity is greater than the weight of *j* with each upstream connected element *i*, we consider the best prima facie cause of *j* to be a spurious cause and we substitute the edge (*i → j*) with the edge (*◊ → j*). Note that in this work we are ignoring deeper discussions of probabilistic causation that aim at distinguishing between actual genuine causes and spurious causes. Instead, we remove spurious causes by using a filter based on correlation because the probability raising of the omnipresent event ◊ is not well defined (see Methods). In addition, we remark that the evaluation for an edge to be a genuine or a spurious cause takes into account all the given events. Because of this, if events are added or removed from the dataset, the same edge can be defined to be genuine or spurious as the set of events included in the model is varied arbitrarily. However, since we do not consider the problem of selecting the set of progression events, we assume that all and only the relevant events for the problem at hand are alreadyknown a priori and included in the model.

Finally, note that this filter is indeed implying a non-negative threshold for the shrinkage-like estimator, when an edge is valid.

**Theorem 1** (Independent progressions). 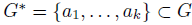 *a set of k* prima facie causes *for some b ∉ G*^*^, *and let* 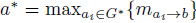. *The reconstructed tree by CAPRESE contains edge ◊ → b instead of a^*^ → b if, for all a_i_ ∈ G^*^*
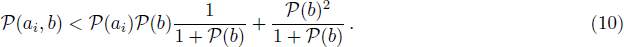

The proof of this theorem can be found in the Supplementary Materials. What this theorem suggests is that, in principle, by examining the level of statistical dependency of each pair of events, it would be possible to determine how many trees compose the reconstructed forest. Furthermore, it suggests that CAPRESE could be defined by first processing the independent progressions filter, and then using *m* to build the independent progression trees in the forest.

To conclude, the algorithm reconstructs a well defined tree (or, more in general, forest).

**Theorem 2** (Algorithm correctness). *CAPRESE reconstructs a well defined tree *𝒯* without disconnected components, transitive connections and cycles*.

Additionally, asymptotically with the number of samples, the reconstructed tree is the correct one.

**Theorem 3** (Asymptotic convergence). *Let T* = (G ∪ {◊},E, ◊) *be the forest to reconstruct from a set of s input samples, given as the input matrix D. If D is strictly sampled from the distribution induced by T and infinite samples are available, i.e., s → ∞ to, CAPRESE with α → 0 correctly reconstructs T.*

The proof of these Theorems are also in the Supplementary Materials. These theorems considered datasets where the observed and theoretical probabilities match, because of *s → ∞* TO. However, data often contains false positives and negatives (i.e., data are noisy) and resistance to their effects is desirable in any inferential technique. With this in mind, we prove a corollary of the theorem analoguos to a result appearing in [22].

**Corollary 1** (Uniform noise). *Let the input matrix D be strictly sampled from the distribution induced by T with sample size s → ∞ to, but let it be corrupted by noise levels of false positives ε+ and false negatives ε_−_. Let* 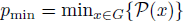, *CAPRESE correctly reconstructs T for α → 0 whenever*

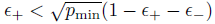
*and ε_+_* ε _−_ < 1.

Essentially, this corollary states that CAPRESE (and so the estimator m) is robust against a noise affecting all samples equally. Also, the fact that it holds for λ → 0 is sound with the theory of shrinkage estimators for which, asymptotically, the corrector factor is not needed to regularize the ill posed problem.

### Optimal Shrinkage-Like Coefficient

Theorem 3 and Corollary 1 state that with infinite samples and mild constraints on the false positive/negative rates we get optimal results with λ → 0. Precisely, for the uniform noise model that we applied to synthetic data (see Methods) we have ε_+_ = ε^→^ = *v*/2, thus the hypothesis required by Corollary 1 is

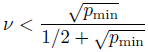

For 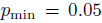, which we set in data generation (see Supplementary Materials), this inequality implies correct reconstruction for v<0.3 (a 15% error rate), with infinite samples. However, we are interested in performance and the optimal value of A in situations in which we have finite sample sizes as well. Here, we empirically estimate the optimal λ value, both in the case of trees and forests, as a function of noise and sample size. In the next section, we assess performance of our algorithm empirically.

In Figure 2, we show the variation of the performance of CAPRESE as a function of λ, for datasets with 150 samples generated from tree topologies. The optimal value, i.e., lowest Tree Edit Distance (TED, see Methods), for noise-free datasets (i.e., *v* = 0) is obtained for λ → 0, whereas for the noisy datasets a series of U-shaped curves suggests a unique optimum value for λ → 1/2, immediately observable for v < 0.15. Identical results are obtained when dealing with forests (not shown here). In addition, further experiments with *n* varying around the typical sample size (*n* = 150) show that the optimal λ is largely insensitive to the dataset size (see Figure 3). Thus we have limited our analysis to datasets with the typical samplesize that is characteristic of data currently available.

Summarizing, Figure 3 and Figure 3 suggest that for sample size below 250 without false positives and negatives the PR raw estimate a suffices to drive reconstruction to good results (TED is 0 with 250 samples), i.e.,

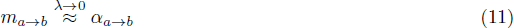

which is obtained by setting λ to a very small value, e.g. 10^−2^, in order to consider at least a small contribution of the correction factor too. Conversely, when *v* > 0, the best performance is obtained by averaging the shrinkage-like effect, i.e.,

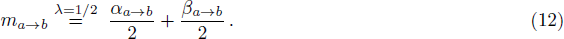

These results suggest that, in general, a unique optimal value for the shrinkage-like coefficient can be determined, even in situations not captured by Theorem 3 and Corollary 1.

## Performance of CAPRESE Compared to Oncotrees

An analogue of Theorem 3 holds for Despers’s oncotrees (Theorem 3.3, [8]), and an analogue of Corollary 1 holds for Szabo’s extension with uniform noise (Reconstruction Theorem 1, [22]). Thus, with infinite samples both approaches reconstruct the correct trees/forests. With finite samples and noise, however, their performance may show different patterns, as speed of convergence may vary. We investigate this issue in the current section.

In Figure 4 we compare the performance of CAPRESE with oncotrees, for the case of noise-free synthetic data with the optimal shrinkage-like coefficient: λ → 0, equation (11). Since Szabo’s algorithm is equivalent to Desper’s without false negatives and positives, we rely solely on Szabo’s implementation[22]. In Figure5 we show an example of reconstructed tree where CAPRESE infers the correct tree while oncotrees mislead a causation relation.

In general, one can observe that the TED of CAPRESE is, on average, always bounded above by the TED of oncotrees, both in the case of trees and forests. For trees, with 50 samples the average TED of CAPRESE is around 6, whereas for Desper’s technique it is around 13. The performance of both algorithms improves as long as the number of samples is increased: CAPRESE has the best performance (i.e., TED ≈ 0) with 250 samples, while oncotrees have TED around 6. When forests are considered, the difference between the performance of the algorithms reduces slightly, but also in this case CAPRESE clearly outperforms oncotrees.

Notice that the improvement due to the increase in the sample size seems to reach a*plateau*, and the initial TED for our estimator seems rather close to the plateau value. This empirical analysis suggests that CAPRESE has already good performances with few samples, a favorable adjoint to Theorem 3. This result has some important practical implications, particularly considering the scarcity of available biological data.

In Figure 6 we extend the comparison to *noisy* datasets. In this case, we used the optimal shrinkage-like coefficient: λ → 1/2, equation (12). The results confirm what observed without false positives and negatives, as CAPRESE outperforms oncotrees up to *v* = 0.15, for all the sizes ofthe sample sets. In the Supplementary Materials we show similar plots for the noise-free case.

We can thus draw the conclusion that our algorithm performs better with finite samples and noise, since less samples are required to get good performances and a higher resistance to false positives and negatives is shown.

## Performance of CAPRESE Compared to Conjunctive Bayesian Networks

Inspired by Desper’s seminal work, Beerenwinkel and others developed methods to estimate the constraints on the order in which mutations accumulate during cancer progression, using a probabilistic graphical model called *Conjuntive Bayesian Networks* (CBN)[14, 15]. While the goal of this research was to reconstruct *direct acyclic graphs* and not trees per se, evidence presented in [24] suggests that, in the absence of noise, these models perform better than oncotrees even at reconstructing *trees*. For this reason, we performed experiments similar to the ones suggested above, comparing CAPRESE to the extension of CBN called *hidden-CBN* (h-CBN) that accounts for noisy genotype observations [15]. This method combines CBNs with a simulated annealing algorithm for structure search and a denoising of the genotypes via the maximum a posteriori estimates to compute the most likely progression. One aspect that complicates a comparison between CAPRESE and (h-)CBN is that the methods assume different models. For example, at the heart of CBN is a monotonicity assumption (i.e., an event can only occur if all its predecessors have occurred) not assumed by CAPRESE. Despite the differences between the model assumptions, we present a preliminary comparison between the methods in Appendix B.3, indicating that we not only outperform oncotrees, but h-CBNs as well. In particular, this suggests that CAPRESE converges much faster than h-CBNs with respect to the sample size, also in the presence of noise.

We also analyze the rate of *false positives/negatives* reconstructed by CAPRESE when (synthetic) data are sampled from DAGs (Appendix B.3). The rate of *false positives* goes to 0 as the sample size increases, implying that CAPRESE is capable of reconstructing a tree subsumed by the underlying causal DAG topology. In addition, the number of *false negatives* approaches a value proportional to the connectivity of the model from which the data was generated. This is expected, since CAPRESE will assign at most one cause to each considered event. However, it should be noted that further experiments with samples selected from a wider array of topologies should be performed to confirm these results and compare both methods in full. While not within the scope of the current paper, these issues will be addressed in future work.

## Case Studies

In the next subsections we apply CAPRESE to real cancer data obtained by *Comparative Genomic Hybridization* (CGH) and *Next Generation Sequencing* (NGS). This shows the potential application of reconstruction techniques to various types of mutational profiles and various cancers.

## Performance on Cancer CGH Datasets

Encouraged by the results in previous sections, we test our reconstruction approach on a real *ovarian cancer* dataset made available within the oncotree package [8]. The data was collected through the public platform SKY/M-FISH [38], used to allow investigators to share molecular cytogenetic data. The data was obtained by using the CGH technique on samples from *papillary serous cystadenocarcinoma* of the ovary. This technique uses fluorescent staining to detect CNV data at the resolution of chromosome arms. While the recent emergence of NGS approaches make the dataset itself rather outdated, the underlying principles remain the same and the dataset provides a valid test-case for our approach. The seven most commonly occurring events are selected from the 87 samples, and the set of events are the following gains and losses on chromosomes arms 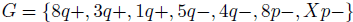 (e.g., 4*q*-denotes a deletion of the q arm of the 4^th^ chromosome).

In Figure 7 we compare the trees reconstructed by the two approaches. Our technique differs from Desper’s by predicting the causal sequence of alterations

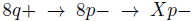
when used either λ → 0 or λ = 1/2. Notice that among the samples in the dataset some are not generated by the distribution induced by the recovered tree, thus comparing the reconstruction for both As is necessary.

At this point, we do not have a biological interpretation for this result. However, we do know that common cancer genes reside in these regions, e.g. the tumor suppressor gene PDGFR on 5*q* and the oncogene MYC on 8*q*), and loss of heterozygosity on the short arm of chromosome 8 is quite common (see, e.g., http://www.genome.jp/kegg/). Recently, evidence has been reported that 8*p* contains many cooperating cancer genes [39].

In order to assign a confidence level to these inferences we applied both parametric and non-parametric *bootstrapping methods* to our results. Essentially, these tests consist of using the reconstructed trees (in the parametric case), or the probability observed in the dataset (in the non-parametric case) to generate new synthetic datasets, and then reconstructs again the progressions (see, e.g., [40] for an overview of these methods and [41] for the use of bootstrap for evalutating the confidence of oncogenetic trees.). The confidence is given by the number of times the trees in Figure 7 are reconstructed from the generated data. A similar approach can be used to estimate the confidence of every edge separately. For oncotrees the *exact tree* is obtained 83 times out of 1000 non-parametric resamples, so its estimated confidence is 8.3%. For CAPRESE the confidence is 8.6%. In the parametric case with false positive and false negative error rates of 0.21 and 0. 027, following [22], the confidence of oncotrees is 17% while the confidence of our method is much higher: 32%. When error rates are forced to 0 the confidence of oncotrees raises to 86.6% and 90.9% respectively.

For the non-parametric case, edges confidence is shown in Table 8. Most notably, our algorithm reconstructs the inference 8*q*+ → 8*p* — with high confidence (confidence 62%, and 26% for 5*q* — → 8*p* —), while the confidence of the edge 8*q+ →>* 5*q*— is only 39%, almost the same as 8*p* — → 8*q*+ (confidence 40%). The confidences are similar with either λ →0 or λ = 1/2.

**Analysis of other CGH datasets**. We report the differences between the reconstructed trees also based on datasets of gastrointestinal and oral cancer ([35, 37] respectively). In the case of gastrointestinal stromal cancer, among the 13 CGH events considered in [35] (gains on 5*p*, 5*q* and 8*q*, losses on 14*q*, 1*p*, 15*q*, 13*q*, 21*q*, 22*q*, 9*p*, 9*q*, 10*q* and 6*q*), oncotrees identify the path progression 
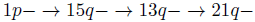
while CAPRESE reconstructs the branch 
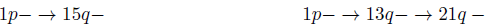

In the case of oral cancer, among the 12 CGH events considered in [37] (gains on 8*q*, 9*q*, 11*q*, 20*q*, 17*p*, 7*p*, 5*p*, 20*p* and 18*p*, losses on 3*p*, 8*p* and 18*q*), the reconstructed trees differ since oncotrees identifies the path

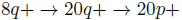

while our algorithm reconstructs the path

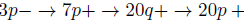

These examples show that CAPRESE provides important differences in the reconstruction compared to oncotrees.

## Performance on Cancer NGS Datasets

In this section we show the application of reconstruction techniques to the validation of a specific relation among recurrent mutations involved in *atypical Chronic Myeloid Leukemia* (ACML).

In [42] Piazza *et al*. used high-throughput *exome sequencing technology* to identity somatically acquired mutations in 64 ACML patients, and found a previously unidentified recurring *missense point mutation* hitting SETBP1. By re-sequencing SETBP1 in samples with A^CML^ and other common human cancers, they found that around 25% of the A^CML^ patients tested positive for SETBP1, while most of the other types of tumors were negative. Assessing the possible relationship between SETBP1 variants and the mutations in many driver ACML oncogenes such as (e.g., ASXLI, TET2, KRAS, etc.) no significant association or mutual exclusion with SETBP1 was found but for ASXLI, which was frequently mutated together with SETBP1, hinting at a potential relation among the events. In particular, ASXLI was presenting either a *non-sense point* or a *indel* type of somatic mutation.

Hence, we reconstructed A^CML^ progression models from the datasets provided in [42], with the goal of assessing a *potential causal dependency* between mutated SETBP1 and ASXLI. A more extensive analysis is postponed, as we only seek to clearly illustrate the functionalities of the algorithm here.

As a first case (Figure 1, left), we treated the ASXLI missense point and indel mutations as indistinguishable, and we merged the two events in the dataset. Afterwards, we separated the two types of mutations for ASXLI (Figure 1, right).

In particular, it is interesting to notice that, when the ASXLI mutations are considered equivalent, the inference suggests that the mutations belong to two independent progression paths (i.e., the independent progression filters “breaks” every potential causal relation among the events). Conversely, when the mutations are kept separate, the progression model suggests that: (i) the missense point mutation hitting SETBP1 can cause a non-sense point mutation in ASXLI and (ii) the observed ASXLI mutations seems to be independent. Concerning edges confidence, as before assessed via non-parametric bootstrap, it is worth noting that the confidence in the indel ASXLI mutation being an early event raises consistently in the latter case.

All in all, it seems that a progression model allows to test the significance of the association firstly observed in [42] and also refines the knowledge by suggesting a specific causal and temporal relations among events. With this this in mind, ad-hoc sequencing experiments might be set up to assess these predictions, eventually providing a strong evidence that could be used to, e.g., synthesize a progression-specific ACML-effective drug.

## Discussion

In this work we have introduced a novel framework for the reconstruction of the causal topologies underlying cumulative progressive phenomena, based on the *probability raising* notion of causation. Besides such a probabilistic notion, we also introduced the use of a *shrinkage-like estimator* to efficiently unravel ambiguous causal relations, often present when data are noisy. As a first step towards the definition of our new framework, we have here presented an effective novel technique called CAPRESE (CAncer PRogression Extraction with Single Edges) for the reconstruction of tree or, more generally, forest models of progression which combines probabilistic causation and a shrinkage-like estimation.

We prove correctness of CAPRESE by showing asymptotic convergence to the correct tree. Under mild constraints on the noise rates, this result holds for the reconstruction problem in the presence of uniform noise as well. Moreover, we also compare our technique to the standard tree reconstruction algorithm based on correlation (i.e., Oncontrees) and to a more general bayesian probabilistic graphical model (i.e., Conjunctive Bayesian Networks), and show that CAPRESE outperforms the state-of-the-art on synthetic data, also exhibiting a noteworthy efficiency with relatively small datasets. Furthermore, we tested our technique on ovarian, gastrointestinal and oral cancer CGH data and NGS leukemia data. The CGH analysis suggested that our approach can infer, with high confidence, novel causal relationships which would remain unpredictable in a correlation-based attack. The NGS analysis allowed validating a causal and temporal relation among key mutations in atypical chronic myeloid leukemia.

One of the strong points of CAPRESE is that it can be applied to genomic data of any kind, even heterogeneous, and at any resolution, as shown. In fact, it simply requires a set of samples in which the presence or the absence of some alterations supposed to be involved in a causal cumulative process have been assessed. Notice also that the results of our technique can be used not only to describe the *progression* of the process, but also to *classify* different progression types. In the case of cancer, for instance, this genome-level classifier could be used to group patients according to the position of the detected individual genomic alterations in the progression model (e.g., at a specific point of the tree) and, consequently, to set up a genome-specific *therapy design* aimed, for instance, at blocking or slowing certain progression paths instead of others, as was studied in [43].

Several future research directions are possible. First, more complex models of progression, e.g. directed acyclic graphs, could be inferred with probability raising and compared to the standard approaches of [14, 15, 44], as we explained in the introduction. These models, rather than trees, could explain the common phenomenon of *preferential progression paths* in the target process via, e.g., *confluence* among events. In the case of cancer, for instance, these models would be certainly more suitable than trees to describe the accumulation of mutations.

Second, the shrinkage-like estimator itself could be improved by introducing, for instance, different correction factors. In addition, an analytical formulation of the optimal shrinkage-like coefficient could be investigated by starting from the hypotheses which apply to our problem setting, along the lines of [45].

Third, advanced statistical techniques such as *bootstrapping* [40] could be used to account for more sophisticated models of noise within data, so as to decipher complex causal dependencies.

Finally, a further development of the framework could involve the introduction of *timed data*, in order to extend our techniques to settings where a temporal information on the samples is available.

## Software Availability

The implementation of CAPRESE is part of the TRanslational ONCOlogy (TRONCO) R package and is available for download at standard R repositories.

## Acknowledgements

We are grateful for the many excellent comments we received from anonymous reviewers.

## B. Supplementary Materials

### B.1 Proofs

Here the proofs of all the propositions and theorems follow.

**Proof of Proposition 1 (Dependency)**

**Figure 2:**
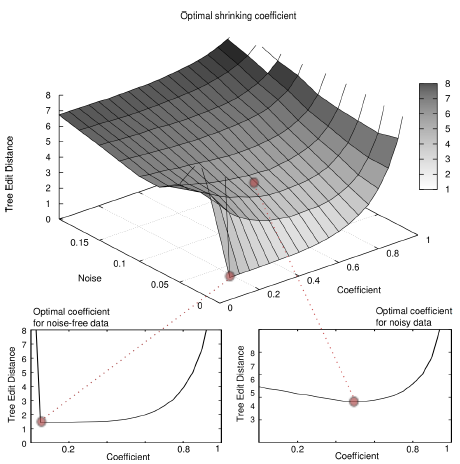
Optimal shrinkage-like coefficient for reconstruction performance. We show here the performance in the reconstruction of trees (TED surface) with *n* = 150 samples as a function of the shrinkage-like coefficient λ. Notice the global optimal performance for λ → 0 when *v* → 0 and for λ ≈ 1/2 when *v* > 0.

**Figure 3:**
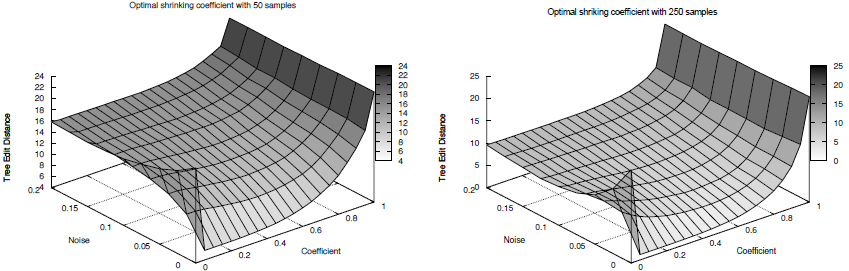
Optimal λ with datasets of different size. We show the analogous of Figure 2 with 50 and 250 samples. The estimation of the optimal shrinkage-like coefficient λ appears to be irrespective of the sample size.

**Figure 4:**
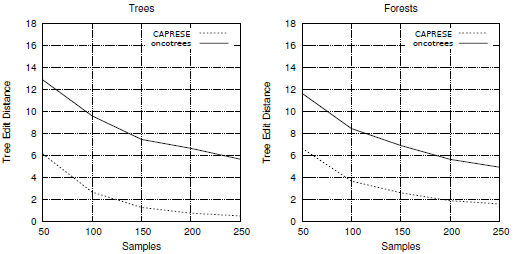
Comparison on noise-free synthetic data. Performance of CAPRESE (dashed line) and oncotrees (full line) in average TED when data are generated by random trees (left) and forests (right). In this case *v* = 0 (no false positives/negatives) and λ → 0 in the estimator *m*.

**Figure 5:**
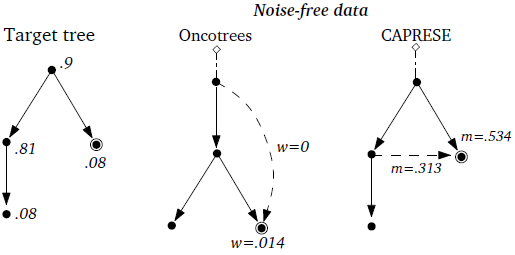
Example of reconstructed trees. Example of reconstruction from a dataset with 100 samples generated by the left tree (the theoretical probabilities are shown, i.e., the doubly-circled event appears in a sample with probability: 08), with *v* = 0. In the sampled dataset oncotrees mislead the cause of the doubly-circled mutation (*w* = 0 for the true edge and *w* = 0:014 for the wrong one). CAPRESE infers the correct cause (the values of the estimator *m* with λ = 1/2 are shown, similar results are obtained for λ → 1).

**Figure 6:**
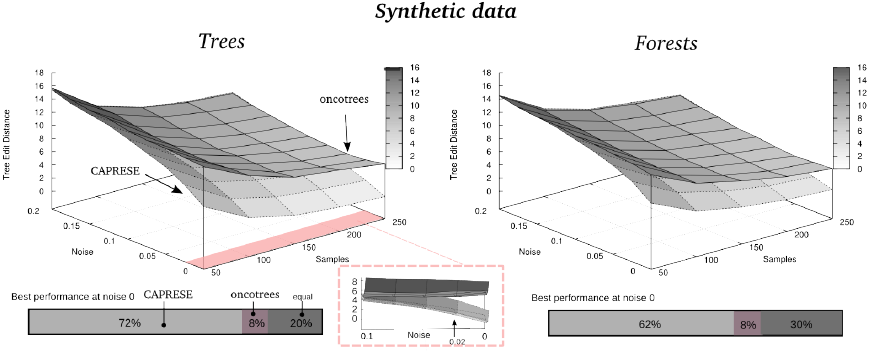
Reconstruction with noisy synthetic data and λ = 1/2. Performance of CAPRESE and oncotrees as a function of the number of samples and noise *v*. According to Figure 2 the shrinkage-like coefficient is set to λ = 1/2. The magnified image shows the convergence to Desper’s performance for*v* ≈ 0:1. The barplot represents the percentage of times the best performance is achieved at *v* = 0.

**Figure 7:**
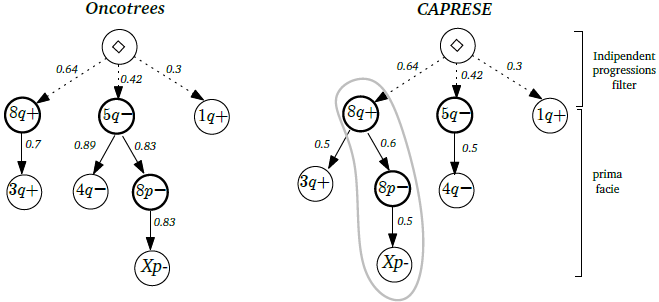
Reconstruction of ovarian cancer progression. Trees reconstructed by oncotrees and CAPRESE (with λ → 0, with λ = 1/2 the same tree is reconstructed). The set of CGH events considered are gains on 8*q*, 3*q* and 1*q* and losses on 5*q*, 4*q*, 8*p* and Xp. Events on chromosomes arms containing the key genes for ovarian cancer are in bolded circles. In the left tree all edge weights are the observed probabilities of events. In the right the full edges are the causation inferred with the pr and the weights represent the scores used by CAPRESE. Weights on dashed lines are as in the left tree.

**Figure 8:**
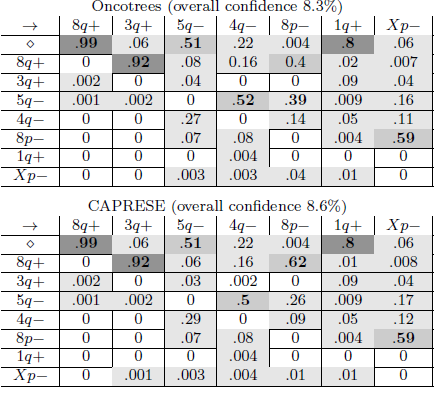
Estimated confidence for ovarian progression. Frequency of edge occurrences in the nonparametric bootstrap test, for the trees shown in Figure 7. Colors represent confidence: light gray is < 0:4, mid gray is in the range [0:4; 0:8] and dark gray is > 0:8. Bold entries are the edges recovered by the algorithms.

**Figure 9:**
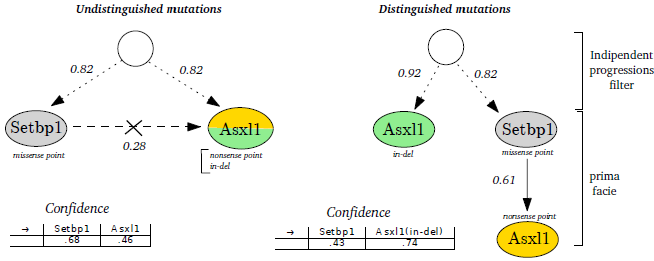
Validating the Setbp1-Asxl1 relation in atypical Chronic Myeloid Leukemia. Progression models where Asxl1 indel and non-sense point are merged (left) and separate (right) suggest that a missense point mutation hitting Setbp1 can cause a non-sense point mutation in Asxl1, that the observed Asxl1 mutations might be independent and that indel Asxl1 is an early event with high confidence.

*Proof*. For ❙ write 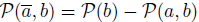, then write the PR as 
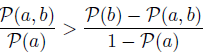
and, since 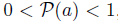, the proposition follows by simple algebraic arrangements of 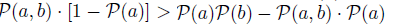. The derivations are analogous but in reverse order for the implication ❘.

**Proof of Proposition 2 (Mutual Probability Raising)**.

*Proof*. The proof follows by Property 1 and the subsequent implication:

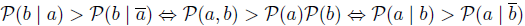

**Proof of Proposition 3 (Probability Raising and Temporal Priority)**

*Proof*. We first prove the forward direction ⟹. Let 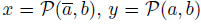 and 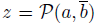. We have two assumptions we will use later on:

1. 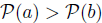 which implies 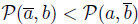, i.e., *x* < *z*.
2. 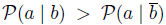 which, when 0 < *x* + *y* < 1, implies by simple algebraic rearrangements the inequality

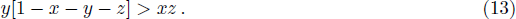

We proceed by rewriting 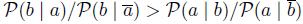 as 
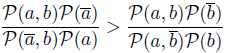

which means that 
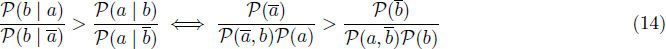

We can rewrite the right-hand side of (14) by using *x, y, z* where 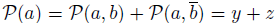 and 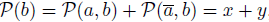, and then do suitable algebraic manipulations. We have 
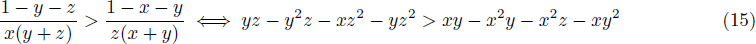

when *x(y* + *z*) ≠0 and *z(x* + *y*) ≠ 0. To check that the right side of (15) holds we show that 
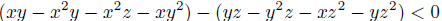

First, we rearrange it to (*x* — *z*)[*y* — *y*^2^ — *xz* — *y*(*x* + *z*)] < 0 so to show that

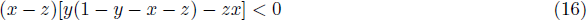

is always negative. By observing that, by assumption 1 we have *z* > *x* and thus (*x* — *z*) < 0, and, by equation (13) we have *y*(1 — *y* — *x* — *z*) — *zx* > 0, we derive

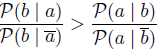

which concludes the ⇒ direction.

The other direction ⇐follows immediately by contraposition: assume that 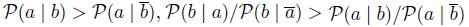 and 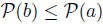. We distinguish two cases:

1. 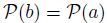 then.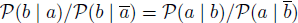
2. 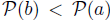, then by symmetry 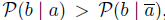, and by the ◹ direction of the proposition it follows that 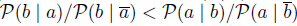.

In both cases we have a contradiction. This completes the proof.

**Proof of Proposition 4 (Monotonic Normalization)**

*Proof*. We prove the forward direction ◹ the converse follows by a similar argument. Let us assume 
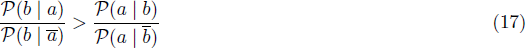
then 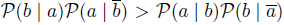. Now, to show the righthand side of the implication, we will show that 
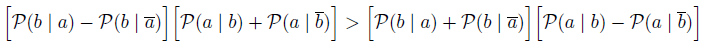

which reduces to show 
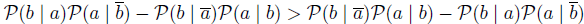

By (17), two equivalent inequalities hold

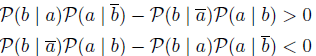
and hence the implication holds.

**Proof of Proposition 5 (Coherence in Dependency- and Temporal Priority)**

*Proof*. We make two assumptions:

1. 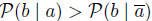which implies α *_a_* → *_b_* > 0.
2. 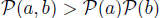
which implies β *_a_* → *_b_* > 0.

The proof for dependency follows by Proposition 1 and its implication:

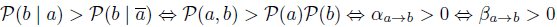

Moreover, being β symmetric by definition, the proof for temporal priority follows directly by Proposition 2 The properties outlined so far are sketched informally in the following diagram.

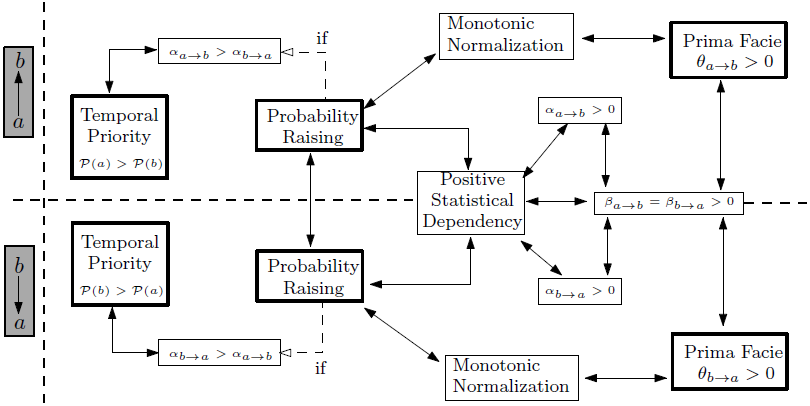

**Proof of Theorem 1 (Independent Progressions)**.

*Proof*. For any *a_i_* ∈ *G*\* it holds that

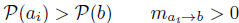

being prima facie, also *a*\* → *b* is the edge selected by CAPRESE being the max{.} over *G*\*. Thus, CAPRESE selects ◊→ *b* instead of *a*^*^ → *b* if, for any *a_i_*, it holds

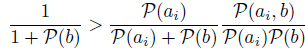

With some algebraic manipulations we rewrite this as

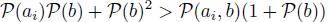

which gives the inequality in the theorem statement.

**Proof of Theorem 2 (Asymptotic convergence)**.

*Proof*. It is clear that CAPRESE does not create disconnected components since, to each node in *G*, a unique parent is attached (either from *G* or ◊). For the same reason, no transitive connections can appear. The absence of cycles results from Properties 3, 4 and 5. Indeed, suppose for contradiction that there is a cycle(*a*_1_,*a*_2_), (*a*_2_,*a*_3_), …, (*a*_n_,*a*_1_)in *E* then by the three propositions we have 
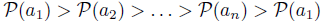
which is a contradiction.

**Proof of Theorem 3 (Asymptotic convergence)**.

*proof*. For any *u* ∊ *G*, when *s* → ∞ the *observed probability P(u)* (evaluated from D) is equivalent to the
product of the probabilities (in *T*) obtained by traversing the forestfrom the root ◊ to *u* (Definition 1).Thus, *P(u)* ∊ (0; 1) since the traversal probabilities are in (0; 1) too, hence all events are distinguishable and (Algorithm 1 reconstructs a tree with the same events set *G* of *T*.

We now observe that the distribution induced by *T* (Definition 1) respects a *single-cause prima facie topology* where to each event is assigned at most a single cause. In other words, Definition 2 holds for any edge (*u; v*) ∊ *E*:

- by the event-persistence property usually assumed in cancer (fixating mutations are present in the progeny of a clone) the occurring times satisfy *t_u_*< *t_v_* which, in a frequentist sense, implies*P(u)* > *P(u)*;

- it holds by construction (Definition 1) that *P(v, u)* = *P(v)*, thus *P(v | u)* = *P(v)*/ *P(u)* which is strictly positive since *P(v)* and *P(u)* are, and that *P(v,ū)* = 0, thus *P(v | u)* = 0.

To correctly reconstruct *T* we rely on the fact that our score *m_u_→_v_* is consistent with the prima facie probabilistic causation because of:

- Proposition 3, which states that *PR* (embedded as α*_u_→ _v_* in *m*) subsumes a good temporal priority model of occurring times, as stated above;

- Proposition 4 and 5 which ensure the monotonicity and sign coherency among α*_u_→_v_* and β*_u_→_v_* in *m*.

Thus, *m* is consistent with a single-cause prima facie topology. We now show that Algorithm 1 reconstructs correctly a generic edge in *E*, and hence also *T*.

Consider anevent *U* ∊ *G* and edge *(u, v)* ∊ *E*. The set of its “candidate” parent events is*G* \ {*v*}, we partition it in three disjoint sets *𝒢*, *𝒮* and *𝒩*:

- *𝒢, genuine:* all the backward-reachable events, in G \ {*v*}, from *v*;

- *𝒮*, *spurious (or ambiguous):* all the events (but v) in the sub-forest which includes the path from ◊ to *v*, which are not in *G*;

- *𝒩*, non prima facie: all other events, i.e., *G* \ ({*v*}∪ *𝒮* ∪ *𝒢*);

Notice that *G* = {*v*}∪ *𝒢*∪ *𝒮*∪ *𝒩*, that *u* ∈ * *𝒢* and that all the effects of *v* are non prima facie to *v* because of the temporal priority, as shown below.

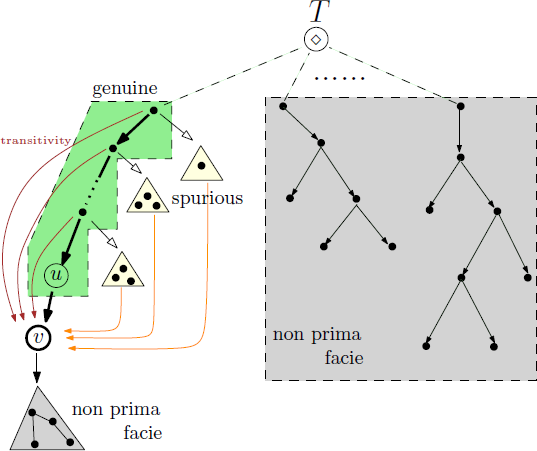

This way of partitioning events according to the structure of *T* subsumes a equivalent partitioning based on the score α ∈ [0,1], which we use to prove correctness of our algorithm: for any *x* ∈ *G* it holds that α*_x_*→*_v_* = 1 if *x* ∈ *G*, 0<α*_x_*→*_v_*<1 if*x* ∈ *S* and α*_x_*→*_v_*<0 if *x* ∈ *N*.We now show that CAPRESE correctly selects *u* ∈ *G*:

- a non prima facie event *x* ∈ *𝒩* either satisfies (Proposition 1)

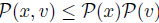

which means that α*_x_*→*_v_* < 0, α*_x_*→*_v_*< 0 (Proposition 5) and thus *m_x_→_v_*< 0 or it is a descendant of *v*,which means that *P(x)*< *P(v)*. By construction, CAPRESE considers as candidate parents of *v* only not descendant events with positive score(see step 3);

- a spurious event *x* * *S* is prima facie to *v* but α_*x*→*v*_;<1 since:

- *𝒫(x) 𝒫(v)*<*𝒫(v,x)*<*P(v)*, otherwise *x* would be backward reachable from *x* and thus in *G*;

- 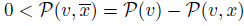which means that 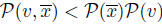

- by all of the above 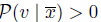which implies that α*_x_*→*_v_*< 1.

Recall now that λ → 0, which means that *m*_*x*→*v*_ ≈ α_*x*→*v*_<1. CAPRESE will thus not select any of these events as cause of *v* if there exist an event with*m_x→v_* = 1, which is actually the case with genuine causes;

- genuine causes are the real cause of *v*, *u*, plus *all* the transitive backward-reachable events. Any *x* of these has maximum score α*_x→v_*=1 since:

- *𝒫(x)𝒫(v)*<*𝒫(v,x)* = *𝒫(v)* and 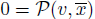;

- by the above 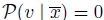 which implies α*_x_*→*_v_* = 1.

Thus, CAPRESE will pick an event from *𝒢*, and not from *𝒮*. We need to show that u is the event with maximum score.

Enumerate the events in*𝒢* as *g*_1_ (which is *u*),…, in a way that

*𝒫(^g1^)*<…<*𝒫(_gk_)*
and recall that this is a *total ordering* induced by the temporal priority, and that this is consistent with coefficient β, which means that
β*_g1_*→*_u_*;> … β*_gk_*→*_u_*.

Thus, in the limit λ → 0

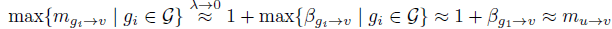

is the event closer in time to *v*, with respect to β. This event, namely *u*, is chosen by the algorithm as the real cause of *v*.

Finally, we show that the last step of the algorithm (the independent progression filter, step 4), does not invalidate the edge *(u,v)*. In fact, the algorithm would replace such an edge with (◊, *v*) if, for all nodes*x*backward-reachable from *v* (i.e., those in *𝒢* ∪ *𝒮*) it was

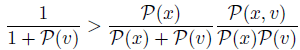

It suffices thus to show that the above inequality is violated just by one of the backward-reachable nodes. We pick just *u* ∈ * *𝒢* and note that

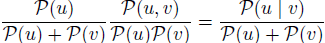

Also, we have that *𝒫(u)*<1, *𝒫(v)*<1 and, by construction, *𝒫*(*u* | *v*)= 1 because all the instances of *v* are co-occurring with those of *u* (but not the converse). Thus, inequality

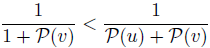

is always true and ensures that edge *(u, v)* is maintained, which concludes the proof.

**Proof of Corollary 1 (Uniform noise)**.

*Proof*. As shown in [22], the uniform rates ∈_+_ and ∈_-_ affect the observed probabilities as follows

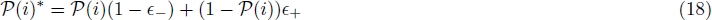

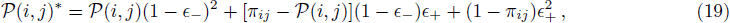

where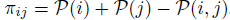 it is imporatantto note (Lemma 1, [22]) that

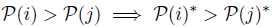

namely uniform noise is still implying temporal priority. Because of this, and since the raw estimate α is monotonic relative to temporal priority, all the derivations for Theorem 3 are still valid in this context, and the algorithm selects the correct genuine cause for each effect.

To guarantee that no valid connection is broken by the independent progressions filter,we again rely on Szabo’s result (Reconstruction Theorem 1, [22]). In particular, for any correctly selected edge *(u, v)* in our algorithm, nsince we implement Desper’s filter (or, analogously, Szabo’s) for independent progressions we do not mistake by deleting *(u,v)* unless also their algorithms do. Since this is not the case when
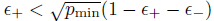 the proof is concluded.

**Figure. 10.**
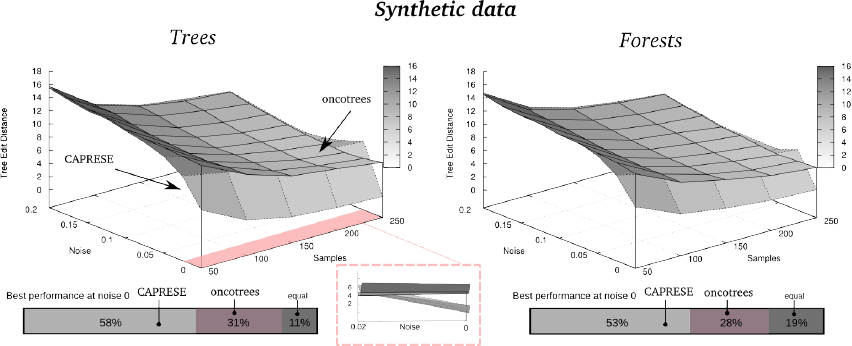
Reconstruction with noisy synthetic data and λ→ 0. The settings of the experiments are the same as those used in Figure 6, but in this case the estimator is shrank by λ→ 0, i.e., λ = 0.01. In the magnified image one can sees that the performance of CAPRESE converges to Desper’s one already for *v* ≈ 0.01, hence largely faster than in the case of λ ≈ 1/2 (Fig. 6.)

### B.2 Synthetic Data Generation

A set of random trees is generated to prepare synthetic tests. Let *n* be the number of considered events and let *p*_min_ = 0.05= 1— *p_max_*, a *single tree* with maximum depth log(*n*) is generated as follows:

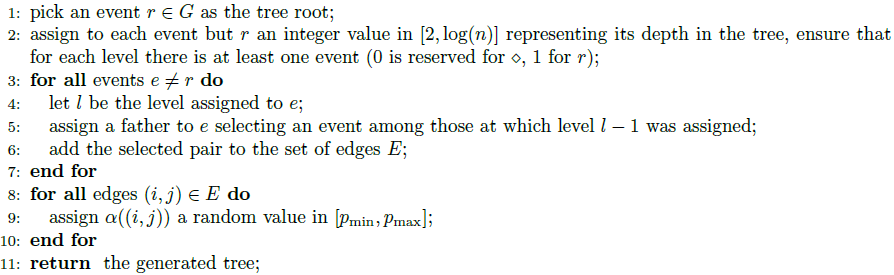

When a forest is to be generated, we repeat the above algorithm to create its constituent trees. These trees (or forests), in turn, are used to sample the input matrix for the reconstruction algorithms, with the parameters described in the main text.

### B.3 Further Results

We show here the results of the experiments discussed but not presented in the main text.

**Reconstruction of noisy synthetic data with** λ → 0. Although we know that λ→ 0 is not the optimal value of the shrinkage-like coefficient for noisy data, we show in Figure 10 the analogue of Figure 6 when the estimator is shrank by λ → 0, i.e., λ = 0.01. When compared to Figure 6 it is clear that a best performance of CAPRESE is obtained with λ≈ 1/2, as suggested by Figure 2.

**Figure. 11.**
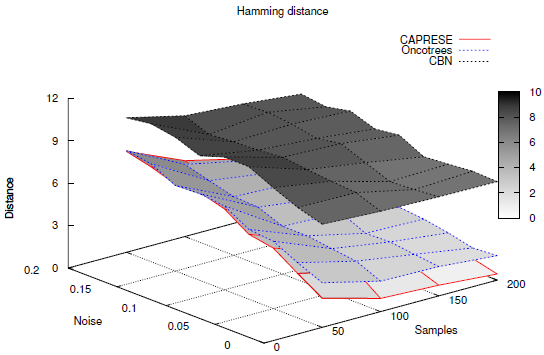
Reconstruction with CAPRESE compared to oncotrees and h-CBN with noisy synthetic data. Performance of CAPRESE compared to oncotrees and h-CBNs as a function of the number of samples and noise *v*. The λ parameter used for CAPRESE is 1/2, and the reconstructed topologies contain 10 nodes each.

**Comparison with hidden Conjunctive Bayesian Networks, h-CBNs**. We here compare the performance of CAPRESE to hidden Conjunctive Bayesian Networks (h-CBN)[15], as well as to oncotrees. The settings of the experiment are slightly different from those of the previous analyses: we used 100 distinct random trees of 10 events each. We ranged the number of samples available for reconstruction from 50 to 200, with a step size of 50. The settings used for running h-CBNs are relatively standard settings: we allowed for 50 annealing iterations with initial temperature equal to 1. Since h-CBNs reconstruct DAGs, it is not possible to quantify its performance using Tree Edit Distance, as we did in the comparison with oncotrees. Instead, we here adopt Hamming Distance (computed on the connection adjacency matrix), as a closely related and computationally feasible alternative for measuring performance [33].

The results of the experiment can be found in Figure 11, and show that CAPRESE clearly outperforms h-CBNs. In particular, it is possible to notice that, for all the analyzed values of noise and sample sizes, both CAPRESE and oncotrees display a (average) Hamming Distance between the reconstructed model and the original tree topology that is significantly lower than h-CBNs, with the largest differences observed in the noise-free case. This result would point at a much faster convergence of CAPRESE with respect to the number of samples, also in presence of moderate levels of noise.

A few remarks are warranted about this experiment. First, in contrast to the comparison with oncotrees, we ran each experiment exactly once rather than averaging the results over 10 repetitions, and on relatively smaller trees. These limitations are a consequence of the extremely high time complexity of the simulated annealing step of h-CBNs. However, the comparison between CAPRESE and h-CBNs shows a so large difference in the performance that we do not expect this to be have significant impact. Second, the results obtained by h-CBNs are perhaps worse than expected based on results in the absence of noise presented in [24], which were however based on a unique tree topology. Yet, this outcome may have been potentially influenced by either the estimation procedure of the noise parameter in h-CBN, the adopted annealing procedure or by the used number of iterations. In future work we plan to extend our algorithm to extract more general topologies and to compare both methods in a greater detail.

**Figure. 12.**
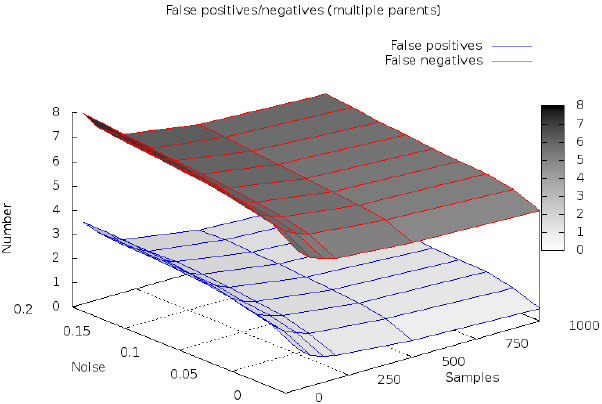
Performance of CAPRESE to reconstruct models with conjunctive parents and noisy data. Performance of CAPRESE measured in terms of the number of *false positives/negatives* in the reconstructed model, when data are generated from directed acyclic graphs with 10 nodes and where each event is caused by at most 3 *conjunctive events* (randomly assigned). The λ parameter is set to 1/2.

**Inference of models with multiple conjunctive parents**. CAPRESE is specifically tailored to reconstruct models with independent progressions and a unique cause for each event (i.e., trees or forests), while other approaches such as CBNs can reconstruct models where multiple conjunctive parents co-occur to cause an effect (i.e., *a ^ b cause c*). It is thus reasonable to use such conjunctive approaches to infer more complex model, in spite of CAPRESE.

However, it is interesting to asses CAPRESE’s performance when (synthetic) data are sampled from a model with multiple parents and noise. By sampling input data from random *directed acyclic graphs* with 10 nodes and where each event is caused by at most 3 conjunctive events (randomly assigned), we assess the number of *false positives* and *false negatives* retrieved in the model reconstructed with CAPRESE. We show the results in Figure 12. Our results indicate that for increasing sample size, the number of false positives approaches 0. Thus, for sufficiently large number of samples, all the causal claims returned by CAPRESE are true. In addition, the number of false negatives is always higher and proportional to the connectivity of the target model. This is to be expected since CAPRESE assigns at most one parent (the cause) to every node.

